# Temperature-responsive contacts in TRPV1 concentrate in the cytoplasmic coupling domains

**DOI:** 10.64898/2026.10.09.757870

**Authors:** Zachary Sottoriva, Davit Potoyan

## Abstract

TRPV1, the heat-activated cation channel of pain-sensing neurons, is a major target for new analgesics, yet where it senses heat remains unresolved. Candidate sensors have been proposed in nearly every part of its structure, from the ankyrin repeats to the pore. Here we ask where the contact network of the closed channel responds to temperature. We simulated human TRPV1 in a multicomponent membrane at six temperatures spanning its activation threshold and three salt concentrations, at atomistic and coarse-grained resolution. We decomposed the contacts that recur across its four identical subunits into collective thermal modes and tested each mode against temperature and, as a control, salinity. Only the leading atomistic mode follows temperature, which explains 59% of its variance and no mode follows salinity at either resolution. The temperature mode places 80% of its variance in the cytoplasmic domains, against 63% of all contact variance, and is most enriched in the linker domain that joins the ankyrin repeats to the transmembrane domain. On heating, the linker releases the C-terminal domain and binds the ankyrin repeats more tightly. Lipid contacts cluster along the transmembrane and TRP domains, and the few that reach the ankyrin repeats and the linker are mostly with the anionic inner-leaflet phosphatidylserine. The coarse-grained model reproduces the atomistic fluctuations and lipid organization, but the temperature mode emerges only at atomistic resolution. Its localization matches that of the dominant temperature mode of the related channel TRPV3. These results narrow the search for the heat sensor. In both channels, the contact network responds to temperature not in the pore but in the cytoplasmic coupling region where the ankyrin repeats, the linker, and the C-terminal domain meet.

**Author summary:** Our sense of burning heat starts with an ion channel, TRPV1, that opens when tissue warms past about 42 degrees Celsius. Where in this large protein heat is detected is still debated. Experiments have implicated almost every region. We used computer simulations to follow the contacts between the channel’s amino acids at six temperatures and three salt concentrations, at two levels of detail. Because each simulation can get stuck in one arrangement, we treated the channel’s four identical subunits as a built-in check and kept only contacts that formed in at least two of them. What remained showed one clear pattern that tracks temperature and none that tracks salt. The pattern sits not in the pore but below the membrane, where the ankyrin repeats, a linker segment, and the tail of the protein meet. As the temperature rises, the linker lets go of the tail and holds the ankyrin repeats more tightly. A simpler, faster model of the channel did not reproduce this pattern. Our results point to specific contacts, between the linker and the tail, that experiments can now test.

## Introduction

Chronic pain remains hard to treat [1]. The analgesics that dominate the market were introduced decades ago [2, 3], and opioids, though effective, interfere with normal brain function and are addictive [3, 4]. Ion channels offer alternative targets [5, 6]. These membrane proteins control the passive flow of selected ions across cell membranes by changing conformation [7, 8] in response to membrane potential, mechanical stimuli, temperature, or acidity [8, 9], and this gating underlies several pain transmission pathways [6].

The transient receptor potential (TRP) superfamily of cation channels is a promising source of pain targets because its members transmit temperature-induced pain sensations [10–13] and take part in signaling pathways linked to disease [14, 15]. Within the superfamily, TRPV1, the founding member of the vanilloid subfamily, is among the most actively pursued targets for treating pain and disease [16–19]. TRPV1 mediates the sensation of noxious heat. It is activated by temperatures above about 42 *^◦^*C and also by endogenous lipid-derived molecules, protons, and capsaicin [20, 21].

Cryo-electron microscopy (cryo-EM) now provides TRPV1 structures detailed enough for simulation, including a recent structure of the human channel [16]. The channel is a homotetramer, and each subunit comprises a transmembrane domain (TMD) and an intracellular domain (ICD) [22] (Fig 1). As in many voltage-gated ion channels, the TMD consists of two modules. The S1–S4 module sits at the channel periphery, and the S5–S6 module forms the central pore together with the pore helix [23–25]. The ICD comprises the N-terminal ankyrin repeat domain (ARD), the linker domain and pre-S1 helix that connect it to S1, the TRP domain that runs along the inner face of the membrane, and the C-terminal domain (CTD) [25–27] (Fig 2).

**Fig 1.**
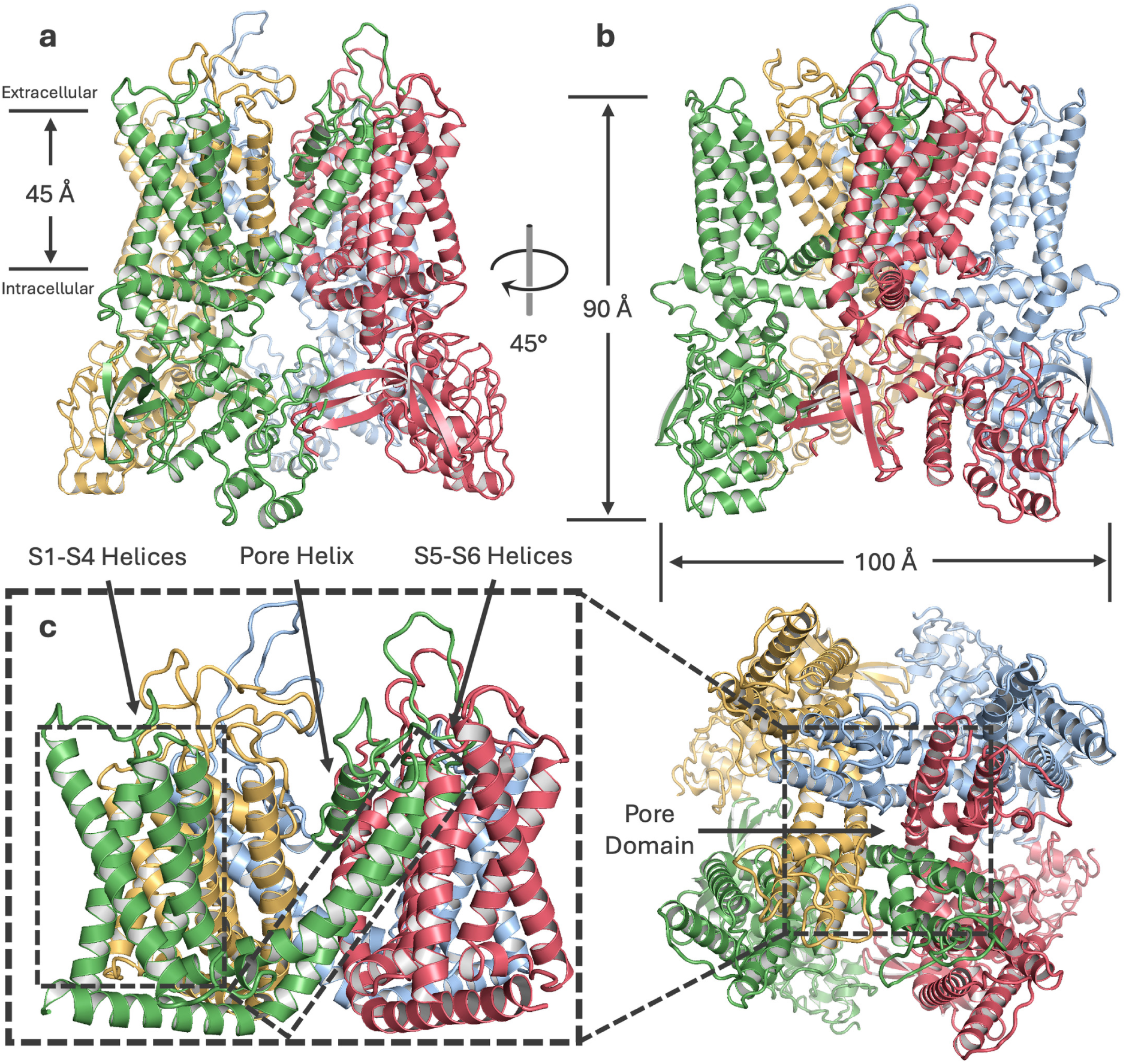
TRPV1 structural overview. a-b. Ribbon views of the tetramer parallel to the membrane, with each subunit in a distinct color. The membrane and the intracellular and extracellular sides are indicated on the left and the overall dimensions on the right. **c** The transmembrane domain viewed perpendicular to the membrane, with selected helices labeled.

**Fig 2.**
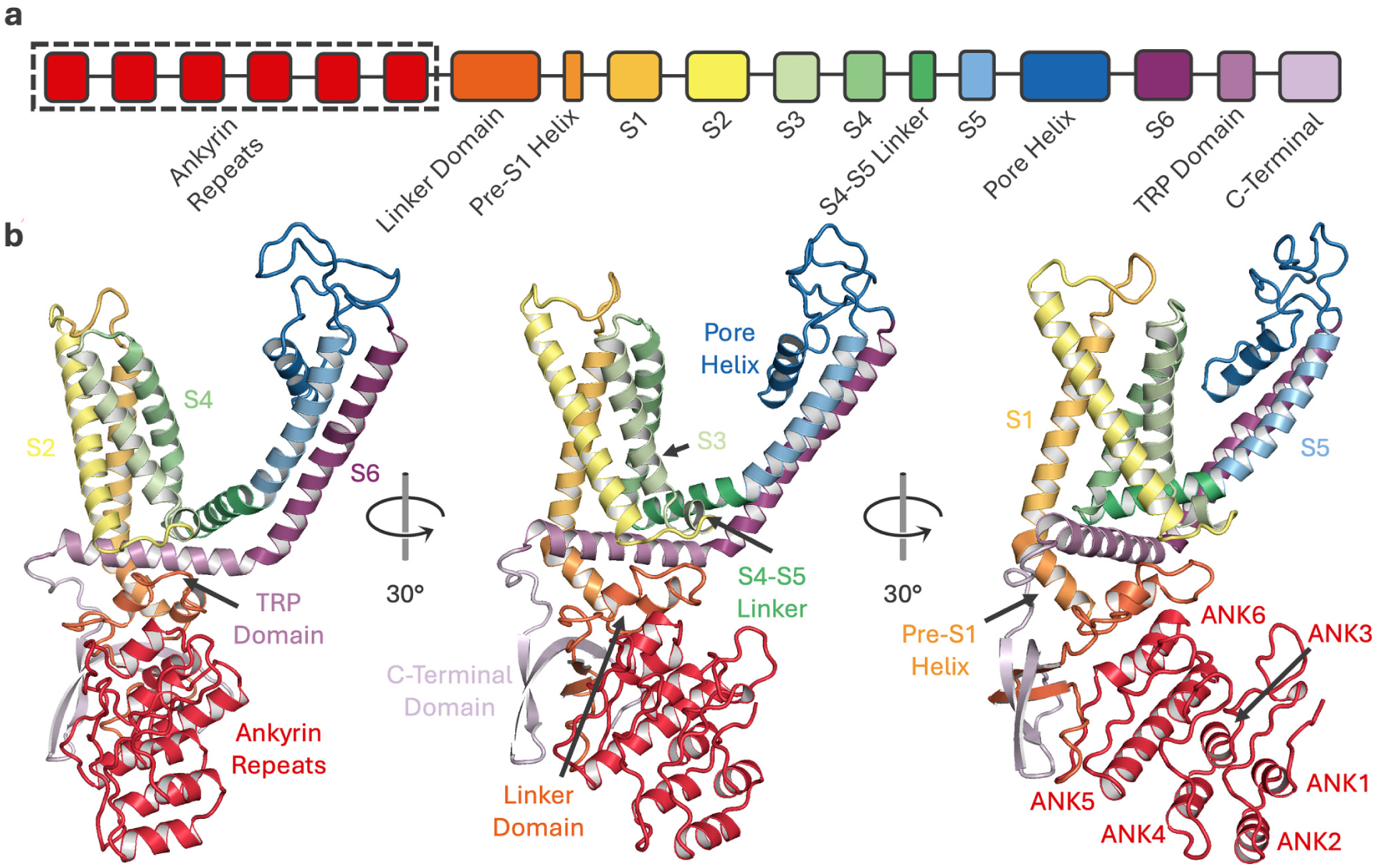
Domain organization of a TRPV1 subunit. **a** Linear diagram of the structural elements; the dashed box marks the six ankyrin repeats of the full-length channel. **b** Ribbon diagrams of one subunit, colored and labeled as in **a**. The resolved structure begins at residue 199, just before ankyrin repeat 3, and contains repeats 3 to 6. This color key is used for the residue bars in Figs 3–7.

The ARD binds other proteins and ATP [27] and modulates channel sensitivity [28]. The C-terminal and TRP domains likewise bind proteins and ligands and contribute to temperature, voltage, and capsaicin sensitivity [29–31]. Heat sensitivity has also been localized to the interface between the TMD and the ICD [32], and further heat-sensing regions have been proposed in the CTD, the outer pore region, and the pore itself [33–39]. Simulations have probed the heat response of the rat channel [22, 40, 41] and, recently, of the human channel [42]. Candidate heat sensors thus span nearly every part of the fold, from the ARD to the pore, leaving open which of them carry the response of the channel to temperature.

Here we ask a narrower question that simulations of the closed channel can address. Where in its structure does the contact network reorganize as temperature rises? We simulated human TRPV1 in a multicomponent membrane at six temperatures from 305 to 330 K, a range that brackets the activation threshold of the channel. Repeating every simulation at three salinities provides a control. TRPV1 operates in extracellular fluid whose ionic strength varies [43, 44], and extracellular cations modulate its activity [43]. Because salt acts on the channel differently from temperature, contacts that follow temperature and not salinity respond to temperature specifically. We ran every simulation and analysis at both atomistic and coarse-grained resolution. The coarse-grained model reaches microseconds at a fraction of the cost, and if it reproduces the atomistic description, it can carry the longer simulations that gating studies of this channel will require. Because one trajectory per condition can trap individual subunits in different conformations, we used the fourfold symmetry of the channel to discard contacts its subunits do not share. The simulations show one collective contact mode that follows temperature and none that follows salinity; the temperature mode appears only at atomistic resolution. It concentrates in the cytoplasmic coupling region formed by the ankyrin repeats, the linker domain, and the CTD. On heating, the linker releases the CTD and binds the ankyrin repeats more tightly.

## Results

We simulated TRPV1 at six temperatures from 305 to 330 K and three salinities, 50, 100, and 150 mM each of KCl and NaCl, which combine the two main cations of neuronal fluids [45]. Each of the 18 conditions was run once at atomistic resolution (400 ns) and once at coarse-grained resolution (4 *µ*s), 36 simulations in all. The runs sample contacts and fluctuations of the closed channel.

### The channel stays folded in every simulation

The fold is preserved in every simulation at both resolutions (S1 and S2 Figs). The RMSD of the C*α* atoms averages 4.23 Å in the atomistic simulations, and that of the backbone beads 5.41 Å in the coarse-grained ones; both are similar across salinities and temperatures. The coarse-grained RMSD fluctuates more, as expected for the smoother energy landscape of Martini [46], even though the elastic network holds the fold of each subunit. These values agree with previous simulations [22, 40, 47]. The radius of gyration of the same atoms averages 44.2 Å (atomistic) and 44.5 Å (coarse-grained) and does not change systematically with temperature or salinity, in agreement with previous work [42, 47]. The channel neither expands nor collapses over the simulated range. The analyses that follow therefore describe an intact channel that stays near its starting, closed conformation.

### Fluctuations peak between structured elements

In the atomistic simulations, the RMSF of the C*α* atoms peaks mostly at borders between structured elements (Fig 3e; Table 1). K241, W273, D301, and K332 fall between ankyrin repeats, consistent with more rigid repeat helices, and the ARD has been implicated in thermal sensitivity [28, 48]. E362, at the ARD/linker border, is also flexible in previous simulations [41, 49]. In the TMD, the peaks fall at the borders between S1 and S2 (K465) [22, 50], S2 and S3 (K504) [22, 49–51], and S4 and the S4–S5 linker (Q560) [41, 49, 50], and at the pore/S6 border (K657) [39, 49, 52, 53], each in a region implicated in gating by temperature, ligands, or protons. E610 also peaks, in an outer-loop region implicated in gating [22, 49, 54], but the loop was rebuilt with Modeller and starts in a different conformation in each subunit (Methods), so the peak may partly reflect the rebuilding. In the ICD, K719 at the TRP/C-terminal border lies at a position implicated in gating [22, 55]. D734 is the exception. It lies within a structured segment, the C-terminal bridge, yet is highly flexible, as in previous simulations [49].

**Fig 3.**
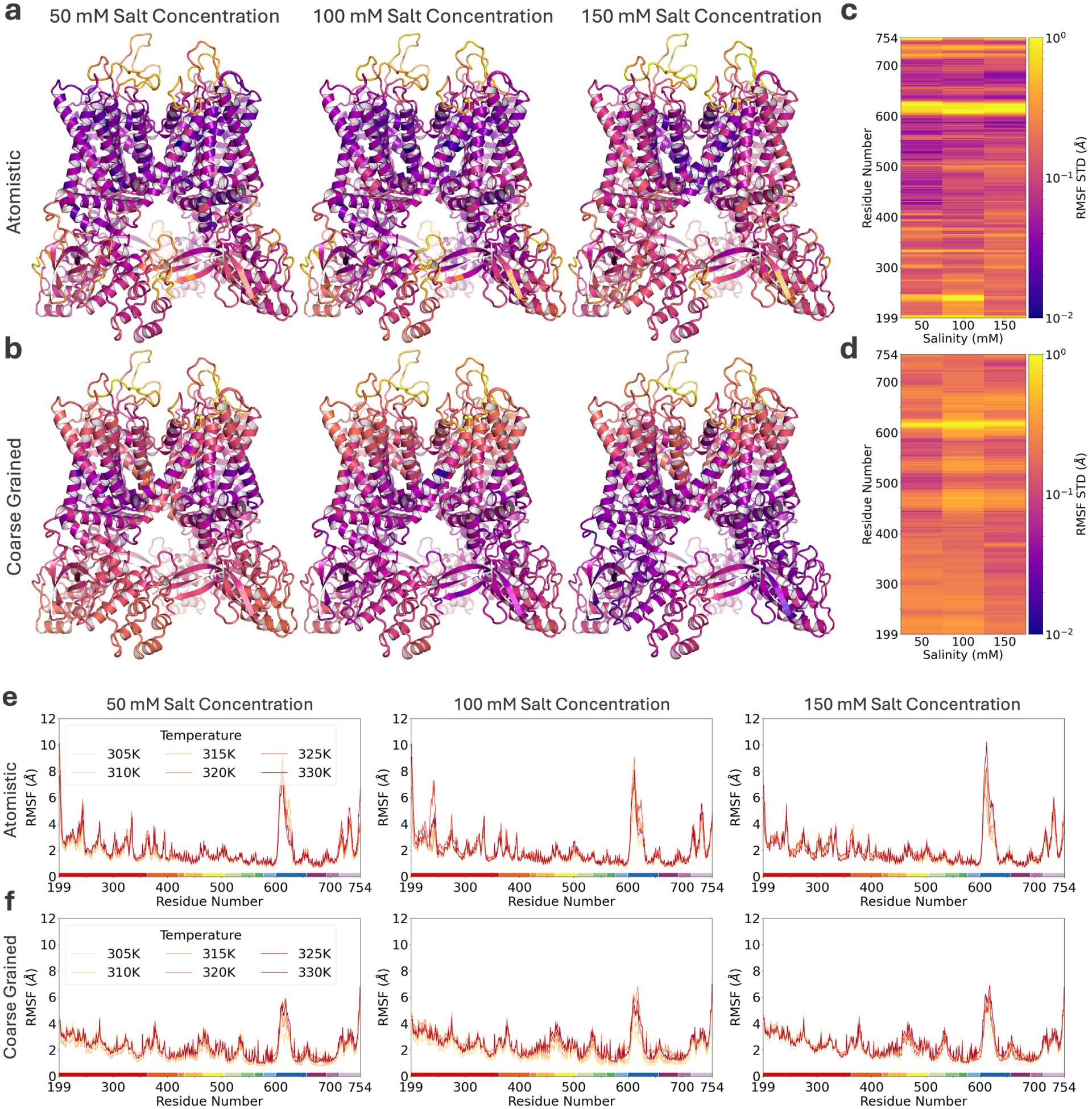
RMSF and its variation with temperature. a-b. Standard deviation of the RMSF over the six temperatures mapped onto the structure for each salinity, atomistic and coarse-grained, colored as in **c-d**. **c-d** The same standard deviation per residue, on a logarithmic scale. **e-f** RMSF per residue at each temperature and salinity, atomistic and coarse-grained. The bar below each residue axis follows the domain key of Fig 2.

**Table 1.**
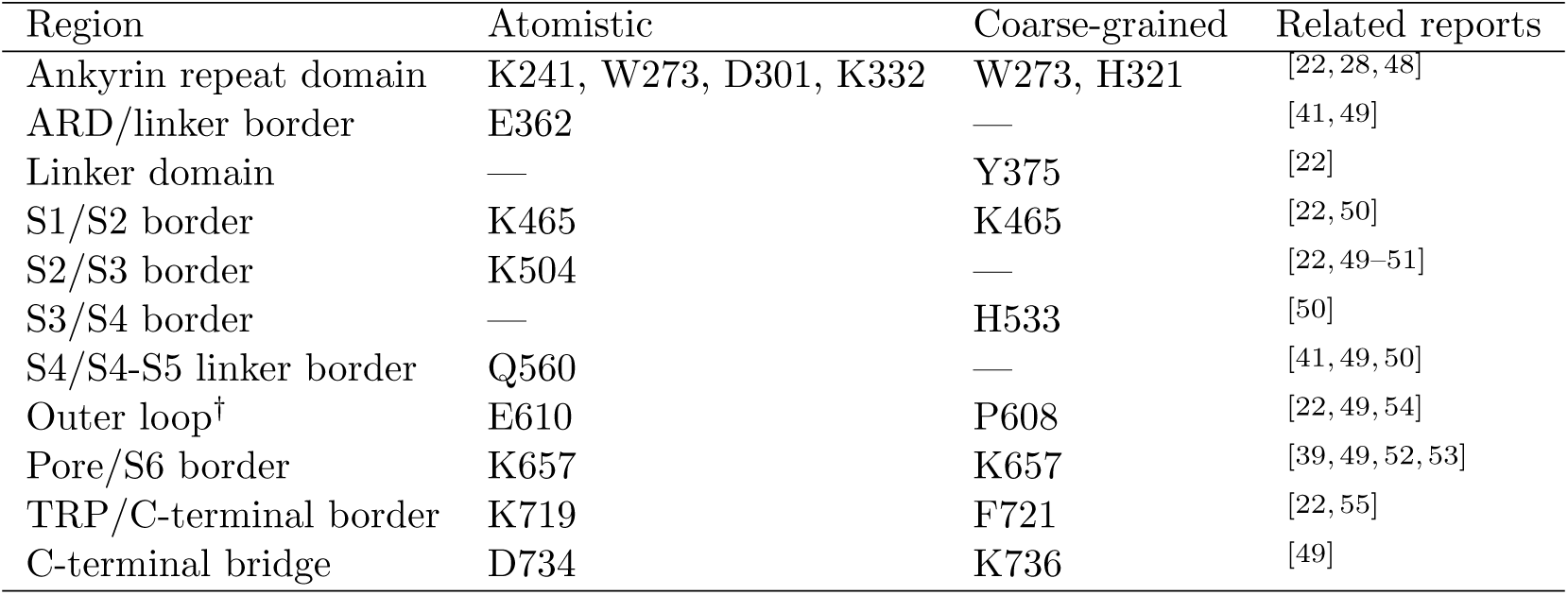
Residues at RMSF peaks.

| Region | Atomistic | Coarse-grained | Related reports |
| --- | --- | --- | --- |
| Ankyrin repeat domain | K241, W273, D301, K332 | W273, H321 | [22, 28, 48] |
| ARD/linker border | E362 | — | [41, 49] |
| Linker domain | — | Y375 | [22] |
| S1/S2 border | K465 | K465 | [22, 50] |
| S2/S3 border | K504 | — | [22, 49–51] |
| S3/S4 border | — | H533 | [50] |
| S4/S4-S5 linker border | Q560 | — | [41, 49, 50] |
| Outer loop<sup>†</sup> | E610 | P608 | [22, 49, 54] |
| Pore/S6 border | K657 | K657 | [39, 49, 52, 53] |
| TRP/C-terminal border | K719 | F721 | [22, 55] |
| C-terminal bridge | D734 | K736 | [49] |

The same residues are those whose RMSF varies most between temperatures (Fig 3a and 3c), and the TMD varies less than the ICD, as expected for a region stabilized by the membrane. Peak heights and the overall spread differ between salinities, but the regions of high RMSF and high spread are the same at all three, so the RMSF shows no effect of salinity on regional fluctuations.

In the coarse-grained simulations, the RMSF of the backbone beads peaks in the ankyrin repeats (W273, H321), in the linker domain (Y375), at the S1/S2 (K465) and S3/S4 (H533) borders, in the outer loop (P608), at the pore/S6 border (K657), at the TRP/C-terminal border (F721), and at the C-terminal bridge (K736) (Fig 3f). Six of these peaks, at W273, K465, P608, K657, F721, and K736, coincide with atomistic peaks at the same position or within a few residues. The two resolutions therefore agree on which regions are most flexible, even after setting aside the two that are flexible by construction, the rebuilt loop and the C-terminal segment near the end of the model. H321 [22], Y375 [22], and H533 [50] lie in regions reported to be temperature sensitive. As in the atomistic simulations, the RMSF shows no effect of salinity on regional fluctuations.

### Symmetry-consistent contacts vary throughout the channel

To see which interactions change with temperature and salinity, we computed the frequency of each residue-residue contact in every simulation, averaged it over the symmetry-equivalent copies of the tetramer, and mapped its variance over the six temperatures at each salinity (Methods).

Before reading these maps, we used the symmetry of the channel to decide which contacts to trust. Because the four subunits are identical, a contact that responds to temperature or salinity should behave alike in all of them, whereas a contact that forms in a single subunit reflects the conformation that subunit happened to adopt. Most contacts are present either in all four subunits or in only one (S8 Fig a), and we kept those present in at least two copies in the median condition (Methods). The filter removed 44% of the atomistic contacts with signal (2,066 of 4,692) and 24% of the coarse-grained ones (825 of 3,369), which carry 32% and 22% of the temperature variance of the raw maps (S3, S5, and S7 Figs). It also removed 84% (atomistic) and 86% (coarse-grained) of the variance of contacts involving the rebuilt 602–626 loop, which starts in a different conformation in each subunit.

The temperature-variance maps at the three salinities share the same regional pattern, with correlations between salinities of 0.68–0.70 (atomistic) and 0.56–0.86 (coarse-grained) (S4 and S6 Figs), and Fig 4 shows the 150 mM maps as representative.

**Fig 4.**
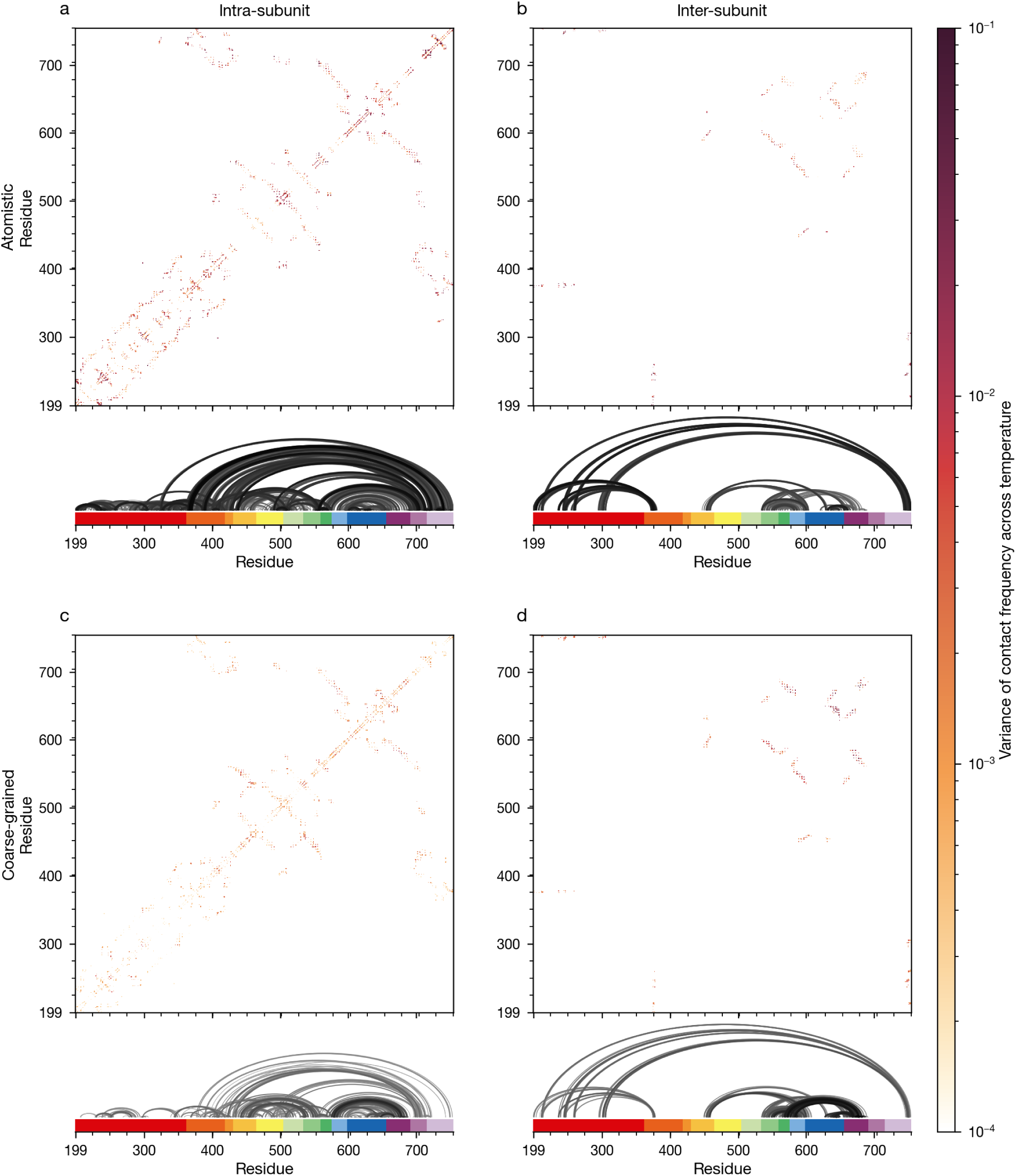
Symmetry-filtered contact variance maps at 150 mM. Variance of each contact frequency across the six temperatures, for symmetry-consistent contacts. **a** Atomistic intra-subunit, **b** atomistic inter-subunit, **c** coarse-grained intra-subunit, and **d** coarse-grained inter-subunit contacts. In each panel, the upper map shows the variance for each residue pair (white where the variance is below 10*^−^*^4^ or the contact is absent), and the lower plot draws an arc for every contact with a variance of at least 10*^−^*^3^, darker and thicker for larger variance, above the domain key of Fig 2. Inter-subunit contacts are plotted at the residue numbers of their two partners. Salinity, 150 mM each of KCl and NaCl; the other salinities are shown in S4 and S6 Figs and the unfiltered maps in S3 Fig.

In the intra-subunit maps (Fig 4a and 4c), contact variance is distributed along the ankyrin repeats, with peaks at the connecting helices, which have been implicated in heat sensitization [21, 32]. Variance also appears between the ankyrin repeats and the linker domain [32] and between the linker and the CTD, which is itself temperature sensitive [20, 33, 34, 56]. Along the TMD it runs from S1 to the TRP domain and is strongest between S4 and the TRP domain, a region whose temperature sensitivity is supported by experiments [21, 36, 50, 57, 58] and simulations [41, 50].

In the inter-subunit maps (Fig 4b and 4d), the largest variances lie between the ankyrin repeats and the linker domain, between the ankyrin repeats and the CTD of the neighboring subunit, which are adjacent in the closed structure [20, 32–34, 56], between S2 and the pore domain [40, 50, 59], and among S4, S5, and S6 [21, 50]. The two resolutions distribute this variance differently. In the atomistic simulations, intra-subunit contacts carry 85% of the filtered temperature variance, whereas in the coarse-grained simulations 58% lies in inter-subunit contacts.

### One contact mode follows temperature and concentrates in the cytoplasmic coupling region

To find contacts that change together and ask whether they follow temperature or salinity, we decomposed the filtered contact frequencies of the 18 simulations by principal component analysis, following the ChACRA framework [60–62]. Each component is a pattern of contacts that strengthen and weaken together, and its score measures how strongly a simulation expresses that pattern. We tested the scores of each component against temperature and salinity over all 18 simulations, with restricted permutations and false discovery rate control across components (Methods; S2 Table).

One component follows temperature, and none follows salinity. The leading atomistic component (15% of the variance) declines with temperature (Fig 5a and 5b). Temperature explains 59% of its variance (*r*^2^ = 0.59, permutation *p* = 0.0007, *q* = 0.013) and salinity 0.3%, and an ANOVA that allows any shape of dependence gives temperature an even larger share (*η*^2^ = 0.80, *p* = 0.006, *q* = 0.10). No other component follows temperature, and none follows salinity (largest *r*^2^ = 0.25, *q* = 0.62). The change concentrates at the two highest temperatures, and the three independent simulations at 330 K have nearly the same score (Fig 5b).

**Fig 5.**
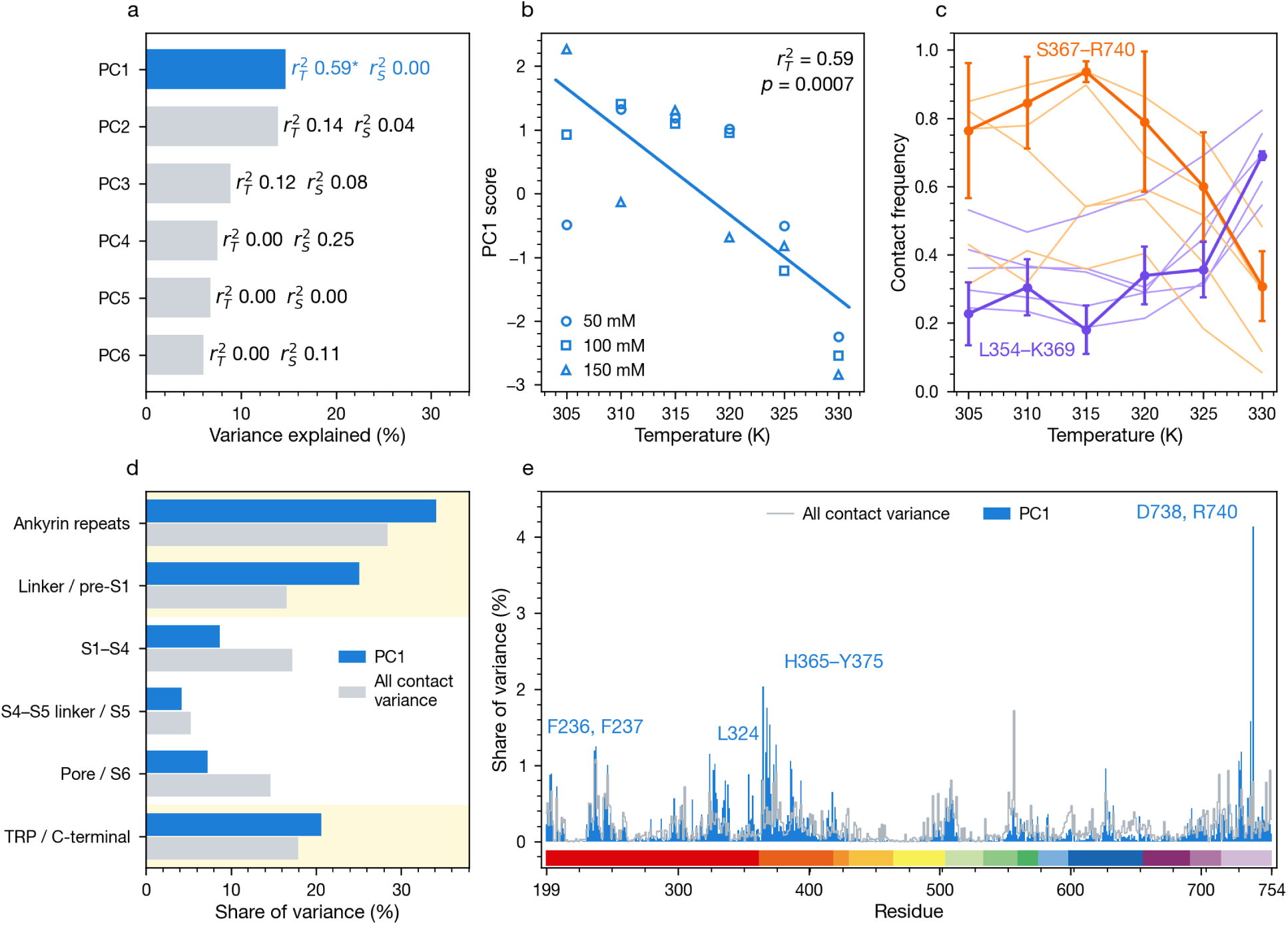
One contact mode follows temperature and concentrates in the cytoplasmic coupling region. Contact-mode analysis of the symmetry-filtered atomistic contacts. **a** The six leading principal components (PCs) by variance explained, with the linear partial coefficients of determination for temperature (*r*^2^_*T*_) and salinity (*r*^2^_*S*_). Blue with an asterisk, temperature mode (q *<* 0.05, restricted permutation test with false discovery rate control); gray, not significant. **b** PC1 score of each of the 18 simulations against temperature; symbols mark the salinity, and the line is the least-squares fit. **c** Frequencies of the twelve contacts with the largest PC1 loadings against temperature, averaged over the three salinities. S367–R740, which weakens on heating, and L354–K369, which strengthens, are drawn with error bars (SD over the three salinities, n = 3 simulations per temperature); the other ten are drawn as thin lines in the color of the labeled contact that changes in the same direction. **d** Share of the PC1 variance in each region next to the share of all contact variance; cytoplasmic regions are shaded. **e** Per-residue share of the PC1 variance along the sequence (bars) and of all contact variance (line), above the domain key of Fig 2.

The temperature mode rearranges the cytoplasmic coupling region (Fig 5c and 5e). Its largest loadings fall into two groups that move in opposite directions. Contacts of R740 in the CTD with linker residues H365, S367, and F370 weaken above 320 K (S367–R740 stays between 0.76 and 0.94 up to 320 K and falls to 0.31 at 330 K). Contacts of the linker with the ankyrin repeats (L354–K369, E327–R368, T336–C387, and E328–S386) strengthen above 320 K as well (L354–K369 stays between 0.18 and 0.36 up to 325 K and rises to 0.69 at 330 K). Above 320 K, the linker thus releases the CTD and binds the ankyrin repeats more tightly. The residues that carry most of the mode are R740, H365, R368, D738, F370, and Y375, followed by F236, F237, and L324 of the ankyrin repeats (Fig 5e).

The mode places 80% of its variance in the cytoplasmic domains, against 63% of all filtered contact variance (Fig 5d). Relative to that baseline, the linker domain and pre-S1 helix carry 1.5 times their share, the ankyrin repeats and the TRP and C-terminal domains 1.2 times, and S1–S4 and the pore and S6 half. Within the TRP and C-terminal bin, the mode sits on the CTD (17% of its variance) rather than on the TRP domain (4%).

Some of the most variable contacts in the raw maps follow neither perturbation. Among them are the contacts of R557, at the junction of S4 and the S4–S5 linker, with E570, T705, and R702; most pass the filter but load mainly on the second component, which tracks neither temperature nor salinity (S2 Table). R557 and E570 line the vanilloid pocket, which held SB-366791 in the cryo-EM structure and is empty in our simulations, so their variability may reflect relaxation of the vacated pocket.

Applying the identical filter and decomposition to the coarse-grained simulations gives a different answer (S8 Fig). No coarse-grained component follows temperature monotonically (largest *r*^2^_*T*_, *q* = 0.32) or salinity, and the leading component (33% of the variance) follows neither. One component, PC3 (12%), depends on temperature without a trend (ANOVA *η*^2^_*T*_, *q* = 0.02; S8 Fig d); it sits in the pore and S6 (52% of its variance, against 42% of all coarse-grained contact variance), around the selectivity filter (M645, L648, Y667). Coarse-grained contact variance is itself concentrated in the TMD (83%, against 37% in the atomistic simulations), and the per-residue variance of the atomistic temperature mode is uncorrelated with that of any of the six leading coarse-grained components (Pearson *r* between *−*0.10 and 0.03). The elastic network that holds the backbone of each coarse-grained subunit, and the restriction of coarse-grained contacts to side-chain beads, change which contacts are free to respond. We therefore base the localization on the atomistic simulations.

The atomistic localization matches the temperature-sensitive contact modes of the related thermo-TRP channel TRPV3, sampled by Hamiltonian replica exchange. The dominant TRPV3 mode clusters on the ARD, and a second clusters around the helix-turn-helix and membrane-proximal domain of the linker region, next to the TRP helix [61]. Two channels sampled by different strategies thus place the temperature response of the contact network in the same part of the fold, the cytoplasmic coupling region where the ankyrin repeats, the linker, and the C-terminus meet, rather than in the pore.

### Lipid contacts cluster along the transmembrane and TRP domains

The membrane around an ion channel can shape its folding and topogenesis [63, 64], function [65, 66], regulation [67, 68], and localization [69, 70]. To see where TRPV1 engages the lipids of the simulated membrane (Fig 6), we analyzed its lipid contacts in the 310 K simulations at the highest salinity.

**Fig 6.**
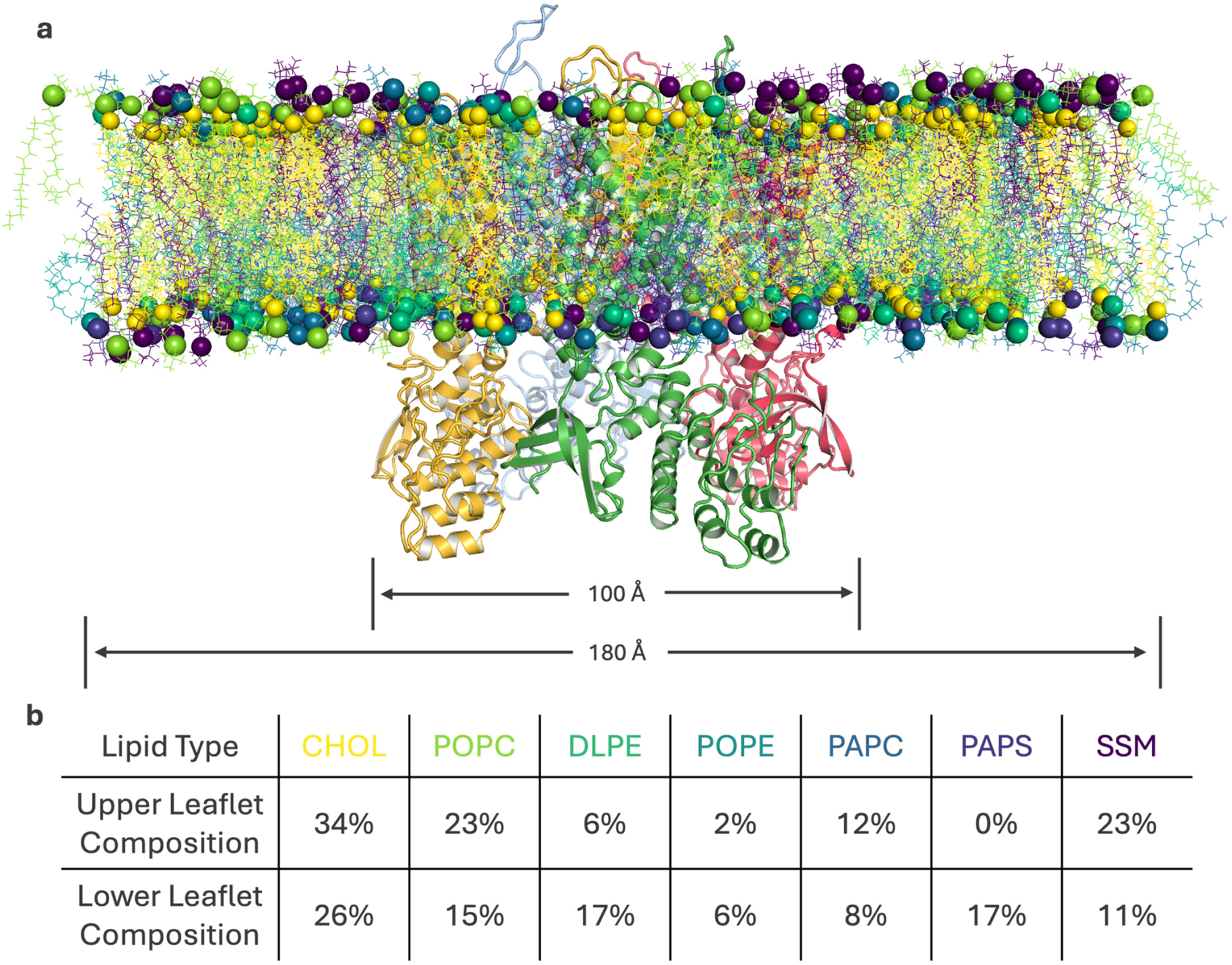
Simulated membrane. **a** TRPV1 embedded in the multicomponent bilayer. Lipids are drawn as a headgroup bead and a tail and colored by species as in **b**; subunits are in distinct colors, and the widths of the protein and the membrane are indicated below. **b** Composition of the outer (upper) and inner (lower) leaflets by lipid species, in percent of lipids.

For every species, the distribution of minimum lipid-protein distances peaks sharply at the first shell, with a tail toward larger distances (Fig 7a and 7b). POPC, PAPC, and PAPS have the narrowest tails, particularly in the coarse-grained simulations, so these species stay in contact with TRPV1 most persistently, consistent with interactions reported for them [71, 72]. The broader tails of the other species indicate that they are more often displaced from direct contact by other lipids.

**Fig 7.**
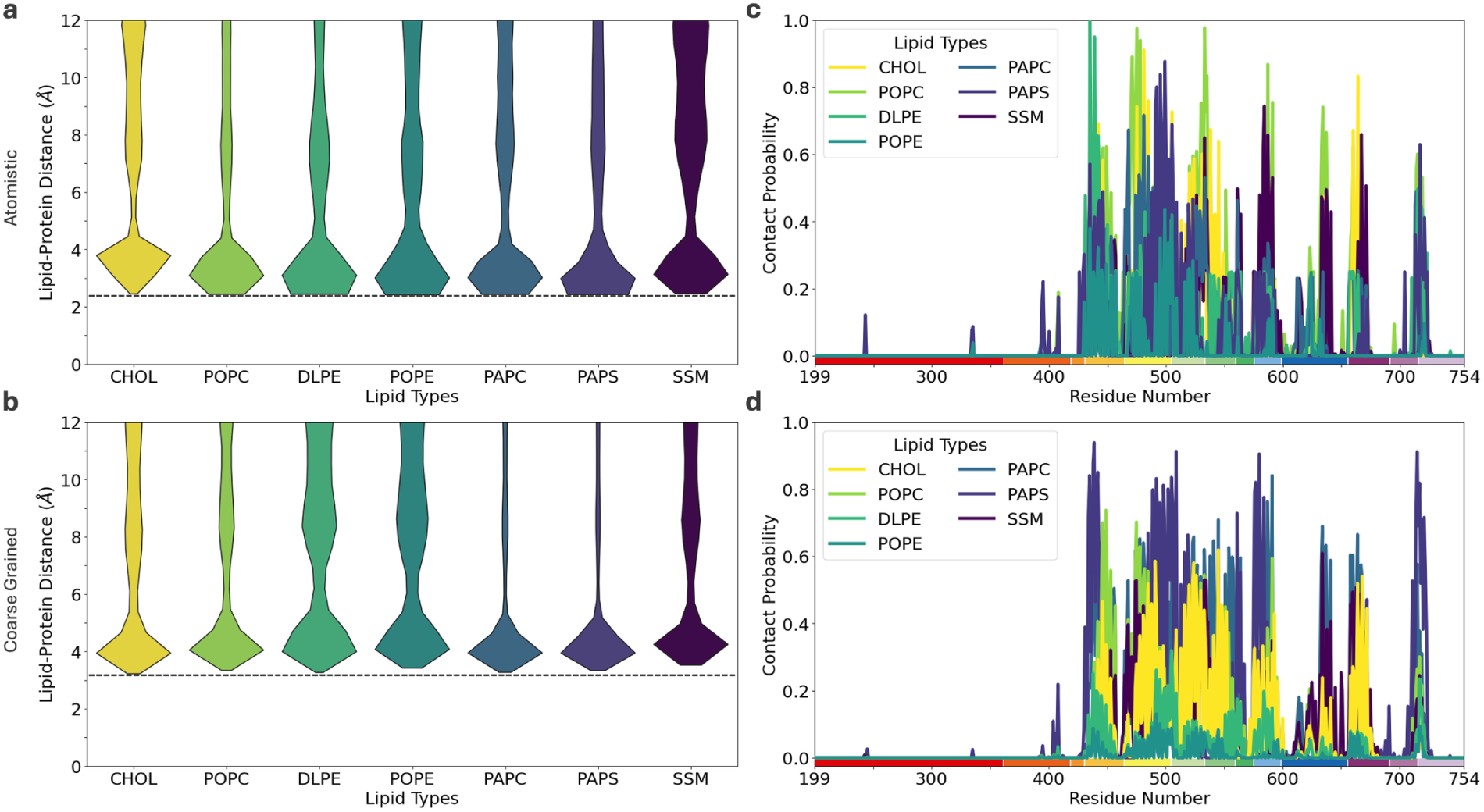
Lipid distances and contacts. a-b. Distributions of the minimum distance between each lipid and the protein, by species, in the atomistic and coarse-grained simulations; colors follow Fig 6, and the dashed line marks the shortest lipid-protein distance observed. The distributions are not normalized for the abundance or leaflet of each species. **c-d** Probability of each residue contacting each lipid species, atomistic and coarse-grained; the bar below the residue axis follows the domain key of Fig 2. Local peak positions are listed in S1 Table. 310 K, 150 mM each of KCl and NaCl.

Lipid contacts form clusters along the sequence (Fig 7c and 7d; S1 Table). Apart from a few isolated peaks, they span S1 to the end of the TRP domain, with clusters at S1, S2–S3, S4, S5, the pore domain, S6, and the end of the TRP domain (L714–A720), separated by the linkers between helices. They thus lie along the TMD, which is embedded in the membrane, and the TRP domain, which lies on its inner face. The pattern is the same at both resolutions, although contact frequencies by species differ. Lipid binding to these regions has been reported for TRPV1 [73] and for the related channel polycystin-2 [74], where specific lipids have been proposed to enable or suppress channel activity. The isolated peaks in the ankyrin repeats and the linker domain are contacts mostly, and occasionally exclusively, with PAPS, at contact probabilities of about 0.1 (atomistic) and 0.02 (coarse-grained). PAPS is the only anionic lipid in the membrane and is confined to the inner leaflet, which faces these domains, so these contacts likely reflect electrostatic attraction of its headgroup to the cytoplasmic domains. Phosphoinositides, absent from our membrane, might occupy the same sites.

## Discussion

Where TRPV1 senses heat has resisted a single answer, and candidate sensors span nearly every part of the channel [75]. Our simulations of the closed human channel address a narrower question, where its network of residue contacts reorganizes as temperature rises, and they point to one region. A single collective mode follows temperature, and it sits below the membrane, in the coupling region where the ankyrin repeats, the linker domain, and the CTD meet. Above 320 K, the linker releases the CTD and binds the ankyrin repeats more tightly. The response is specific to heat, because salinity, the control, leaves it unchanged, and it is reproducible. The three salinity series act as independent replicates of the temperature scan, the contacts of the mode recur across the four subunits, and the three simulations at 330 K, one at each salinity, reach nearly the same score (Fig 5b).

This location agrees with experiments that give the cytoplasmic domains a thermal role. Chimeras show that the N-terminal region connecting the ankyrin repeats to the first transmembrane segment, where our mode is most enriched, is crucial for heat sensing in vanilloid receptor channels [32]. The isolated ankyrin repeat domain changes structure at temperatures that activate the channel [48], the CTD is itself temperature sensitive [33, 34], and atomistic simulations have proposed thermal roles for the C-terminal and adjacent linker domains [49]. Cryo-EM structures of the rat channel at a range of temperatures show that heat opens the capsaicin-bound channel in steps, with rearrangements across several domains before the outer pore moves, and they implicate a network of interacting residues in heat sensing [21]. A contact network that reorganizes in the closed channel, below the membrane and away from the pore, fits the first of these steps.

Other studies place thermal sensitivity in the membrane domains. The isolated S1–S4 domain undergoes a temperature-dependent transition [50], and the pore turret sets species-specific activation thresholds [58]. In the closed channel, S1–S4 and the pore carry half their share of the temperature mode, although the outer pore is among the most mobile parts of the structure. Mobility and coordinated response are different properties, and only the coordinated reorganization is a candidate for the coupling that gating requires. One reading, consistent with the stepwise opening seen by cryo-EM [21], is that heat reorganizes the coupling region first and engages the membrane domains as the channel moves toward opening. Earlier simulations, run without the cytoplasmic domains [40] or far above the threshold, at up to 72 *^◦^*C [22] and 400 K [42], emphasized the pore domain and the S2–S3 and S4–S5 linkers. The closest comparison to our design, contact modes of TRPV3 sampled by replica exchange, places the temperature response in the same cytoplasmic region [61].

The comparison of resolutions adds a lesson for modeling. Martini 3 [76] reproduces the atomistic fluctuations and lipid organization of this channel but not its temperature response, and the elastic network that holds each subunit, together with contacts restricted to side-chain beads, limits which interactions can respond. Contact-level questions about heat sensing therefore need atomistic detail for now.

Lipids add a further layer. Protein-lipid contacts cluster along the transmembrane and TRP domains, away from most of the temperature mode, but the few that reach the ankyrin repeats and the linker, mostly with the anionic inner-leaflet PAPS, fall where the mode is concentrated. Phosphoinositides, which regulate TRPV1 through distinct binding sites [73] and are absent from our membrane, might occupy the same positions. These positions mark where to test whether anionic lipids tune the response of the coupling region to heat.

The results make specific predictions. If the rearrangement lies on the gating pathway, mutating R740 or its linker partners should shift the temperature sensitivity of the channel. Simulations started from an open structure would show whether the rearrangement between the linker and the CTD differs between the closed and open states. The answer matters beyond mechanism. TRPV1 antagonists have yet to become medicines, and the hyperthermia they cause remains poorly understood [75]. A heat response concentrated away from the pore and the vanilloid pocket marks a region where heat sensing might be modulated more selectively. Of the many regions proposed as heat sensors, the closed channel concentrates the temperature response of its contact network in one, the cytoplasmic coupling region where the ankyrin repeats, the linker domain, and the C-terminal domain meet.

## Materials and methods

### Protein model

We built the model from the cryo-EM structure of human TRPV1 in complex with the antagonist SB-366791 (PDB 8GFA) [16], which at 2.29 Å is the highest-resolution human structure in that study (the apo structures 8GF9 and 8GF8 are at 2.58 and 2.9 Å). We removed the ligand, which occupies the vanilloid site, and all other non-protein molecules, including 36 resolved POPC lipids. The model spans residues 199–754 of each subunit (of 839), the range resolved in 8GFA, from just before ankyrin repeat 3 through the CTD, and both termini were left charged. It lacks the distal C-terminus, which binds calmodulin and phosphatidylinositol 4,5-bisphosphate (PIP_2_), and its charged C-terminal residue, N754, lies 14 residues from R740. Residues G602–S626 of the outer pore turret, unresolved in 8GFA, were rebuilt with Modeller [77] through ChimeraX [78]. Because the loop was built independently in each subunit, the four copies start in different conformations. After superposition of the subunit cores, which agree to 0.03 Å, the loops differ by 16–18 Å C*α* RMSD.

### Membrane and solution

We embedded the protein in a 180 *×* 180 Å bilayer of 953 lipids whose asymmetric composition follows simplified models of biological membranes [79, 80] (Fig 6). The bilayer contains cholesterol (CHOL); the phosphatidylcholines POPC (1-palmitoyl-2-oleoyl) and PAPC (1-palmitoyl-2-arachidonoyl); the phosphatidylethanolamines DLPE (1,2-dilauroyl) and POPE; stearoyl sphingomyelin (SSM); and the anionic phosphatidylserine PAPS (1-palmitoyl-2-arachidonoyl), which is confined to the inner leaflet. The membrane contains no phosphatidylinositol or PIP_2_. Each system contained equal concentrations of KCl and NaCl, 50, 100, or 150 mM of each salt, together with 64 Na^+^ ions that neutralize the net charge of the protein and the PAPS lipids. Total monovalent salt was therefore 100, 200, or 300 mM, plus about 27 mM of neutralizing Na^+^, and the K^+^:Na^+^ ratio rose from 0.65 to 0.85 across the three salinities. We refer to the conditions by the concentration of each salt.

### Atomistic simulations

Systems were built with CHARMM-GUI [81–92], solvated and ionized with GROMACS [93], and simulated with GROMACS using the CHARMM36m force field [94] and the TIP3P water model [95]. Each system contained about 540,000 atoms, including about 132,000 water molecules. After steepest-descent minimization, we equilibrated each system in five steps, two at constant volume (125 ps each, 1 fs time step) and three at constant pressure (250 ps at 1 fs, then two of 500 ps at 2 fs), relaxing in stages the position restraints on the protein backbone (from 4000 to 500 kJ mol*^−^*^1^ nm*^−^*^2^), the side chains, and the lipids, and the dihedral restraints. Production runs lasted 400 ns with a 2 fs time step, at 305, 310, 315, 320, 325, and 330 K for each salinity. Temperature was controlled with the v-rescale thermostat [96] (*τ* = 1 ps; protein, membrane, and solvent coupled separately) and pressure with the semi-isotropic Parrinello–Rahman barostat [97] at 1 bar (*τ* = 5 ps; compressibility 4.5 *×* 10*^−^*^5^ bar*^−^*^1^). Electrostatics used particle-mesh Ewald summation [98] with a 1.2 nm real-space cutoff. Lennard-Jones interactions were cut off at 1.2 nm, with a force switch from 1.0 nm during minimization and the first two equilibration steps and a potential-shift modifier during the remaining equilibration and production; the CHARMM36m protocol prescribes the force switch throughout. All bonds were constrained with LINCS [99] (order 4, one iteration), whereas the CHARMM36m protocol constrains only bonds to hydrogen.

### Coarse-grained simulations

We converted the protein to the Martini 3 representation [76] with Martinize2 [100] (vermouth 0.12.0), which maps up to four heavy atoms onto each bead, assigns secondary structure with DSSP, and adds an elastic network on the backbone beads of each subunit (force constant 700 kJ mol*^−^*^1^ nm*^−^*^2^, cutoff 0.9 nm; 2,330 bonds per subunit and none between subunits). Insane [101] built a bilayer of the same size and composition (970 lipids, with 85 rather than 80 PAPS and Martini DPSM for sphingomyelin) and added the solvent. Ions were added with GROMACS as in the atomistic systems, with 69 Na^+^ neutralizing the net charge, for about 51,000 beads in all. After steepest-descent minimization, we equilibrated each system in five constant-pressure steps with the semi-isotropic Berendsen barostat [102] (time steps of 2, 5, 10, 15, and 20 fs; 4.75 ns in total), relaxing the restraints on the protein backbone and the lipid headgroups. Production runs lasted 4 *µ*s with a 20 fs time step, at the same six temperatures and three salinities, with the v-rescale thermostat [96] and the semi-isotropic Parrinello–Rahman barostat [97] (1 bar, *τ* = 12 ps, compressibility 3 *×* 10*^−^*^4^ bar*^−^*^1^). Electrostatics used the reaction-field method [103] (*ɛ_r_* = 15), and electrostatic and Lennard-Jones interactions were cut off at 1.1 nm with a potential shift. Constraints used LINCS [99] of order 8 with two iterations, which avoids artificial temperature gradients in Martini bilayers [104].

### Stability analysis

We computed the root mean square deviation (RMSD) of the C*α* atoms (atomistic) or backbone beads (coarse-grained) from the first production frame with gmx rms, the radius of gyration (*R_g_*) with gmx gyrate, and the per-residue root mean square fluctuation (RMSF) with gmx rmsf, each over the full production run. We report the variation of the RMSF with temperature as its standard deviation over the six temperatures at each salinity.

### Contact analysis

All contact analyses (the raw and filtered variance maps and the contact-mode analysis) used every fifth frame (100 ps atomistic, 1 ns coarse-grained) after the first 50 ns of each atomistic and the first 100 ns of each coarse-grained run, about 3,500 and 3,900 frames per simulation. By then the RMSD has risen to near its plateau (S1 Fig). Two residues are in contact in a frame when any pair of their heavy atoms lies within 4.5 Å (atomistic) or any pair of their beads within 6.0 A° (coarse-grained); residues adjacent in sequence are excluded. Coarse-grained contacts were restricted to pairs involving at least one side-chain bead, because the elastic network fixes the backbone beads. The frequency of a contact is the fraction of frames in which it is present. Coarse-grained residues, numbered 1–556 in the Martini topology, were shifted by 198 to the 8GFA numbering used throughout.

Because TRPV1 is a C4 homotetramer, each contact has four symmetry-equivalent copies, one per subunit or subunit interface, or two for contacts across the channel diagonal. We averaged each contact over its expected copies, a copy that never forms counting as zero; these averages are the raw contact frequencies. To remove contacts that form in only some subunits, we counted, in each condition in which the strongest copy of a contact exceeded a frequency of 0.2, how many copies exceeded both 0.1 and 30% of the strongest one, and we retained the contact if at least two copies were present in the median such condition (both copies for diagonal contacts). The retained contacts form the filtered set. Contact variance maps show, for each salinity, the variance of each contact frequency over the six temperatures. Simulation parameter files and analysis code are available at https://github.com/zamydm/GIMLET.

### Contact-mode analysis

To find contacts that change together, we applied principal component analysis to the 18 *× N* matrix of filtered contact frequencies (mean-centered and unscaled), following the ChACRA framework (chemically accurate contact response analysis) [60–62] and using MDAnalysis [105] for trajectory processing. The 18 conditions yield at most 17 informative components. The eigenvalue permutation test of ChACRA retained 12 atomistic and 10 coarse-grained components and therefore does not discriminate here. We instead tested the scores of each component directly against the two perturbations. The tests use all 18 simulations rather than averages, because a straight line through three salinity averages fits with *R*^2^ *≥* 0.5 for half of all random components. In the balanced 6 *×* 3 design the temperature and salinity regressors are orthogonal, so the partial coefficient of determination of temperature in the joint linear model, *r*^2^, equals the squared correlation of the 18 scores with temperature; *r*^2^ is defined likewise for salinity. P-values come from 10^5^ restricted permutations, with temperature labels shuffled within each salinity and salinity labels within each temperature, and the false discovery rate was controlled across the 17 components separately for each factor (Benjamini–Hochberg). A component is a temperature or salinity mode when its q-value is below 0.05. As a check that allows any shape of dependence, we also report an additive two-way analysis of variance (ANOVA) with the same permutations, whose *η*^2^ is the share of a component’s variance explained by a factor (S2 Table).

To locate a component, we assigned the variance it carries in each contact (its eigenvalue times the squared loading) half to the region of each residue and summed within six regions that follow the domain key of Fig 2. The regions are the ARD (199–361), the linker domain and pre-S1 helix (362–430), S1–S4 (431–559), the S4–S5 linker and S5 (560–598), the pore and S6 (599–691), and the TRP domain and CTD (692–754). The same sum over all components gives the share of all contact variance in each region, the baseline against which the localization of a component is judged. Per-residue profiles are the same sums for each residue.

### Lipid analysis

We analyzed protein-lipid contacts in the 310 K simulations at 150 mM of each salt. For each lipid we computed the minimum distance between its heavy atoms and those of the protein (atomistic) or between its beads and the protein beads (coarse-grained), and we plotted the distribution of these distances by species. A residue contacts a lipid when any of its heavy atoms lies within 4.5 Å of a lipid heavy atom (atomistic) or any of its beads within 6.0 Å of a lipid bead (coarse-grained); we report the probability of each residue contacting each lipid species.

## Supporting information

**S1 Fig. Simulation RMSD. a** RMSD of the C*α* atoms in the atomistic simulations over 400 ns. Colors correspond to temperature, and the panels show 50, 100, and 150 mM of each salt from left to right. **b** RMSD of the backbone beads in the coarse-grained simulations over 4 *µ*s, presented as in **a**.

**S3 Fig. Raw contact variance maps at 150 mM.** Variance of each contact frequency across the six temperatures before the symmetry filter, presented as in Fig 4. **a** Atomistic intra-subunit, **b** atomistic inter-subunit, **c** coarse-grained intra-subunit, and **d** coarse-grained inter-subunit contacts. The frames and contact definitions are those of Fig 4, so the difference between the two figures is the filter alone.

**S4 Fig. Symmetry-filtered intra-subunit contact variance maps. a** Atomistic and **b** coarse-grained simulations, at 50, 100, and 150 mM of each salt from left to right. Maps and arcs are presented as in Fig 4.

**S5 Fig. Raw intra-subunit contact variance maps.** As S4 Fig, before the symmetry filter.

**S6 Fig. Symmetry-filtered inter-subunit contact variance maps. a** Atomistic and **b** coarse-grained simulations, at 50, 100, and 150 mM of each salt from left to right. Maps and arcs are presented as in Fig 4.

**S7 Fig. Raw inter-subunit contact variance maps.** As S6 Fig, before the symmetry filter.

**S8 Fig. Symmetry filter and coarse-grained contact modes. a** Median number of symmetry copies present for four-copy contacts with signal, atomistic and coarse-grained; contacts with fewer than two copies, left of the dashed line, were removed. Half-integer values arise from medians over an even number of conditions. **b** Principal components of the filtered coarse-grained contacts by variance explained, with *r*^2^ and *r*^2^ as in Fig 5a. No component is significant by the trend test; the dagger marks the component whose temperature dependence is significant by the shape-free ANOVA (q *<* 0.05). **c-d** Scores of coarse-grained PC2 and PC3 for each simulation against temperature; symbols mark the salinity, and the dashed line joins the means over salinities. **e** Variance share of the atomistic temperature mode (PC1) and of coarse-grained PC3 in each region, relative to the share of all contact variance at the same resolution (dashed line, equal to the baseline); cytoplasmic regions are shaded.

**S1 Table. Local peaks of residue-lipid contact probability.** Local peaks of the residue-lipid contact probabilities in Fig 7, grouped by residue proximity, for the atomistic (left) and coarse-grained (right) simulations. An empty cell indicates no local peak.

**S2 Table. Temperature and salinity dependence of the leading contact modes.** For the six leading principal components at each resolution, the columns give the variance explained; the linear partial coefficients of determination for temperature (*r*^2^_*T*_) and salinity (*r*^2^_*S*_), with Benjamini–Hochberg q-values from restricted permutations, which set the mode labels; and the effect sizes *η*^2^ of the shape-free two-way ANOVA with their q-values. The q-values control the false discovery rate across the 17 components of each resolution, separately for each factor. n.s., not significant.

## Acknowledgments

The authors acknowledge fruitful discussions with Divyanshu Shukla and Dan Burns.

---

**S2 Fig.**
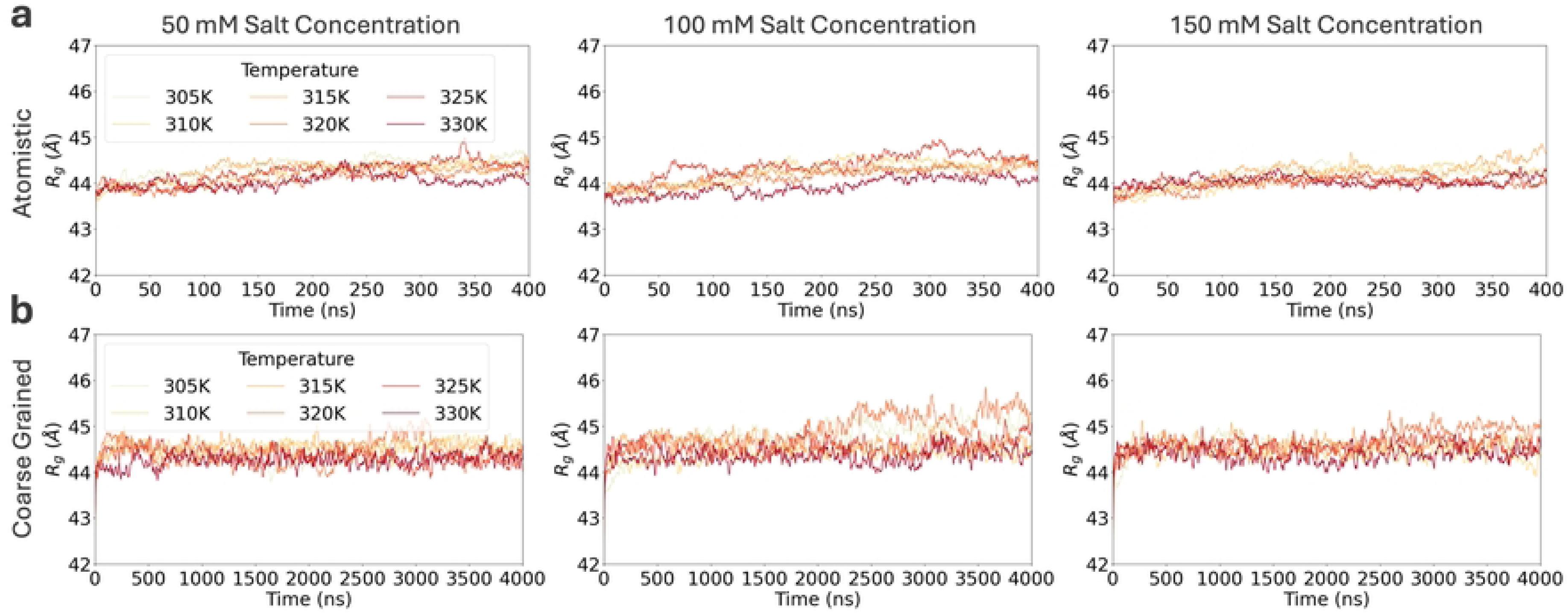
Radius of gyration. **a** *R_g_* of the C*α* atoms in the atomistic simulations over 400 ns. Colors correspond to temperature, and the panels show 50, 100, and 150 mM of each salt from left to right. **b** *R_g_* of the backbone beads in the coarse-grained simulations over 4 *µ*s, presented as in **a**.

